# Broadly reactive influenza antibodies are not limited by germinal center competition with high affinity antibodies

**DOI:** 10.1101/2020.05.11.089953

**Authors:** Rachael Keating, Jenny L. Johnson, David C. Brice, Jocelyn G. Labombarde, Alexander L. Dent, Maureen A. McGargill

## Abstract

Enhancing the generation of broadly reactive influenza antibodies is a pertinent goal towards developing a universal influenza vaccine. While antibodies that bind conserved influenza epitopes have been identified in humans, the frequency of these antibodies is typically very low. The predominant theory is that antibodies specific for conserved influenza epitopes are limited in germinal centers by competition with high affinity antibodies specific for the variable region of the virus. Here, we show that reducing germinal center formation and removing competition with high affinity antibodies was not sufficient to increase broadly reactive influenza antibodies or enhance protection against distinct influenza subtypes. These data disprove the prevailing hypothesis that broadly reactive influenza antibodies are rare due to competition in germinal centers. Additionally, levels of IgM antibodies specific for the variable region of HA persisted in mice in the absence of germinal centers, further demonstrating that immunodominance can be established independent of germinal centers. Our data also highlight the protective capacity of germinal center-independent IgM antibodies, which are not typically considered when testing correlates of protection, and offer an alternate target for delivering a universal influenza vaccine.

**IMPORTANCE:** It is estimated that 250,000 – 650,000 individuals worldwide die each year from seasonal influenza infections. Current vaccines provide little protection against newly emerging strains. Thus, considerable effort is focused on enhancing the generation of broadly reactive influenza antibodies in order to develop a universal influenza vaccine. However, broadly reactive antibodies are rare and the factors that limit their generation are not completely understood. Our data disprove the prevailing hypothesis that broadly reactive influenza antibodies are rare due to completion in the germinal centers with antibodies to the variable HA head. Understanding the factors limiting antibodies specific for conserved regions of the influenza virus is imperative for developing a universal vaccine, which could potentially circumvent a global pandemic

## Introduction

Seasonal influenza vaccines do not effectively protect against novel strains that emerge each year. Consequently, there is considerable interest in developing a universal influenza vaccine that induces immunity to epitopes conserved across different virus subtypes, thereby providing long-lasting, heterosubtypic protection against multiple influenza strains. The antibody response to current influenza vaccines is dominated by antibodies specific for the globular head domain of the surface protein, hemagglutinin (HA) (1). Antibodies targeting the HA head neutralize the virus by preventing the virus from binding host epithelial cells. To escape immune detection, the influenza virus mutates key residues of the HA head region, which frequently gives rise to novel strains. Thus, the most effective antibodies generated by current vaccines are specific for the most variable region of the virus, and therefore only provide strain-specific protection. In addition to the variable HA head, broadly reactive influenza antibodies specific for conserved epitopes of influenza, including those in the membrane proximal stalk region of HA, have been identified in humans (2–5). Broadly reactive influenza antibodies have the capacity to protect against infection with multiple influenza subtypes and therefore, form the basis of a universal vaccine. However, these ‘broad spectrum’ B cell clones are extremely rare and consequently, do not contribute significantly to the antibody response following vaccination (1, 6–9). Indeed, it is estimated that antibodies specific for the HA head are 1000-fold more prevalent than antibodies targeting conserved epitopes (10). A better understanding of factors limiting antibodies specific for conserved regions of the influenza virus is critical for developing a universal vaccine, which could potentially circumvent a global pandemic.

Following influenza infection or vaccination, naïve B cells encounter antigen in the draining lymph nodes or the spleen. Influenza-specific B cells are activated and can differentiate into short-lived plasma cells, memory B cells, or seed germinal centers (11). B cells with the highest avidity for antigen are selected to develop in germinal centers through a series of interactions with T follicular helper cells (T_FH_), which provide essential factors that promote proliferation and further hypermutation of B cells. As T_FH_ are limiting, BCRs with the highest affinity have a competitive advantage over lower affinity BCRs (11). Following affinity maturation in the germinal center, B cells can differentiate into antibody-secreting plasma cells or memory B cells.

Although it is well established that the B cell response following influenza infection or vaccination is heavily biased toward epitopes in the variable head region of HA relative to conserved HA stalk epitopes, the mechanisms that mediate this immunodominance are not completely understood (12). Several factors contribute to the scarcity of broadly reactive antibodies, including steric hindrance of conserved epitopes, antigen quantity, naïve B cell precursor frequency, B cell receptor avidity, and immunization route (11–14). Furthermore, some antibodies specific for conserved epitopes on the HA stem exhibit polyreactivity, which may promote deletion or anergy of B cells generating these antibodies to circumvent autoimmunity (15, 16). The prevailing hypothesis for the paucity of broadly reactive influenza antibodies is that antibodies specific for the conserved epitopes are out-competed in germinal centers by high avidity antibodies specific for the HA head region (11, 12). This is supported by the finding that immunization with the conserved HA stalk region, in the absence of the variable HA head, can elicit a robust antibody response (14, 17). Additionally, mice treated with a low dose of rapamycin during influenza infection had impaired germinal center formation, reduced influenza-specific IgG, and an unexpected increase in broadly reactive antibodies that protected mice against subsequent heterosubtypic infections (18). These data suggest that BCR avidity and competition in germinal centers are major factors limiting development of antibodies specific for conserved influenza epitopes. Moreover, these findings imply that reducing high avidity antibodies specific for the variable HA epitopes could permit the expansion of broadly reactive influenza antibodies, and consequently increase heterosubtypic immunity. Therefore, we tested whether inhibiting germinal center formation was sufficient to limit high avidity antibodies and increase production of broadly reactive influenza antibodies. We found that blocking germinal center formation impaired the synthesis of high avidity IgG antibodies. However, removing competition by high avidity antibodies was not sufficient to increase the prevalence of antibodies specific for the conserved influenza epitopes, or increase heterosubtypic protection following secondary infection. Importantly, we also discovered that heterosubtypic protection correlated with an increase in influenza-specific IgM antibodies. These data demonstrate that B cell competition in germinal centers is not the main determinant responsible for immunodominance or for the paucity of broadly reactive influenza antibodies elicited against conserved epitopes.

## Results and Discussion

### Inhibition of germinal centers reduces influenza-specific IgG, but not IgM antibodies

To test the hypothesis that the antibody response to conserved regions of influenza is limited by competition with high avidity antibodies in germinal centers, we utilized two different methods to reduce germinal center formation and thwart synthesis of high avidity IgG antibodies. In the first model, germinal centers were blocked by administering anti-CD40L blocking antibody on days 6 and 8 following infection (19, 20). For the second model, germinal centers were reduced by conditional deletion of *Bcl-6* mediated by CD4-Cre (*Bcl6*^*fl/fl*^.*CD4-Cre; Bcl6*^*CD4*^). It was previously demonstrated that *Bcl6*^*CD4*^ mice have normal B cell and T cell development, but lack T_FH_ and germinal centers following immunization (21). Importantly, the effector T cell response to immunization is largely intact in *Bcl-6*^*CD4*^ mice. In both models, mice were infected intraperitoneally (i.p.) with an H3N2 strain of influenza (A/HK/x31 (HKx31)). When given i.p., the influenza virus undergoes limited replication, yet produces the full spectrum of proteins to mimic a vaccination (22, 23). We previously showed that mice treated with a low dose of rapamycin, during influenza infection, had reduced numbers of germinal centers and an increase in broadly reactive influenza antibodies, which protected mice from subsequent heterosubtypic infections (18). Therefore, we also treated mice daily with rapamycin (or PBS as a control) to compare whether germinal center formation was reduced to a similar extent by anti-CD40L or elimination of T_FH_ cells.

To assess the extent of germinal center formation, the mediastinal lymph nodes (MLN) that drain the peritoneal cavity were removed 15 days after vaccination and analyzed for germinal centers by immunohistochemistry staining for Bcl-6. Anti-CD40L blockade reduced germinal center formation to a level comparable to daily rapamycin treatment (Fig. 1, A and B). Similarly, *Bcl6*^*CD4*^ mice had a profound defect in germinal center formation compared to wildtype controls (Fig. 1, C and D). Although germinal center numbers were lower in *Bcl6*^*CD4*^ mice compared to mice treated with rapamycin or anti-CD40L, B cells expressing Bcl-6 were still identifiable, but not organized into germinal centers (Fig. 1C). Together, these data show reduced germinal center formation following influenza infection via three distinct mechanisms: anti-CD40L treatment, T_FH_ elimination, and rapamycin treatment.

**Fig. 1.**
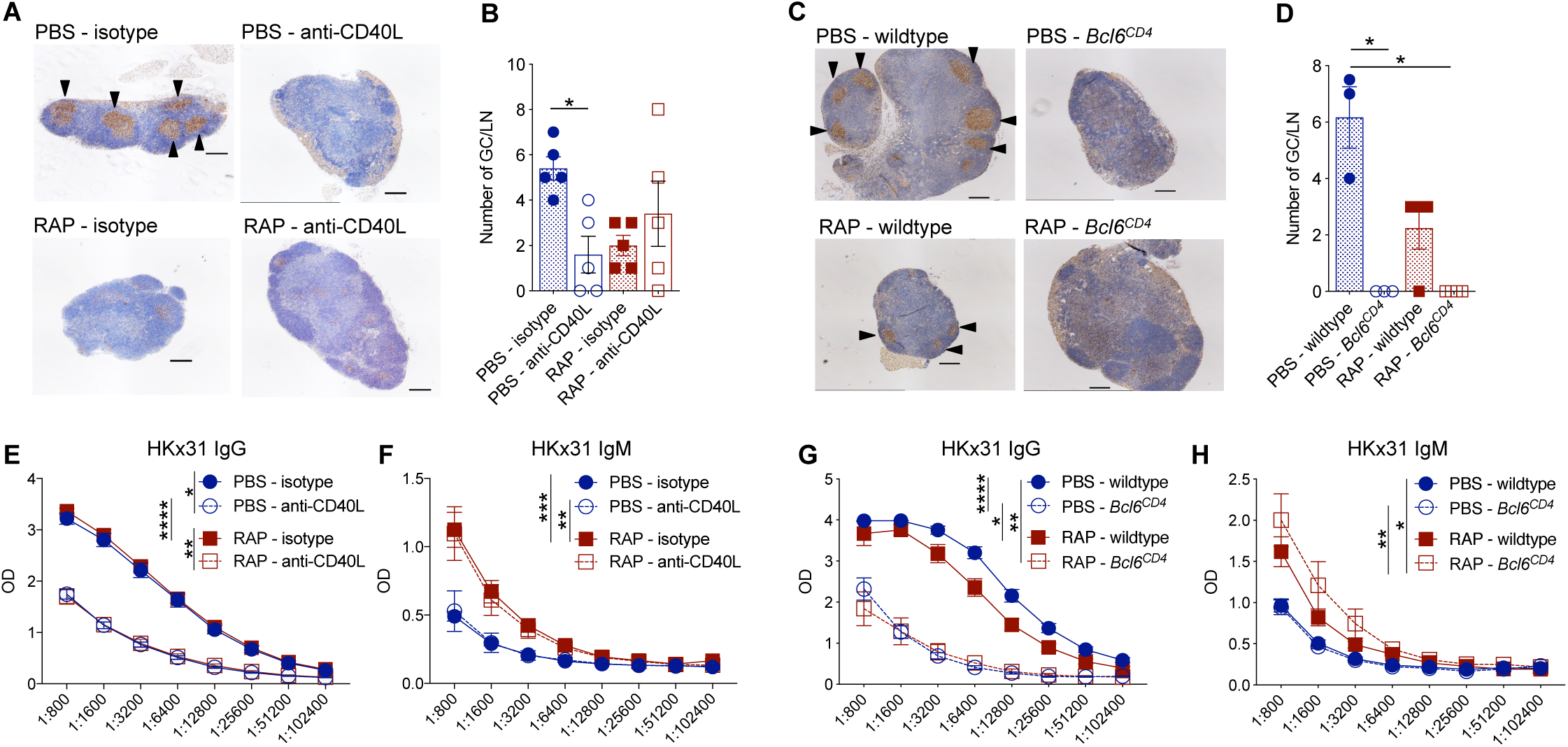
Inhibition of germinal centers reduces influenza-specific IgG, but not IgM antibodies. **(A)** C57BL/6 mice were infected i.p. with HKx31 and treated daily with rapamycin or PBS as a control. Mice were also given anti-CD40L or isotype control, 6 and 8 days following immunization. Mediastinal lymph nodes were removed 15 days after immunization and sections were stained with anti-Bcl6 to identify germinal centers (indicated by arrow heads). **(B)** The average number of germinal centers per lymph node is depicted. Data are representative of two independent experiments. Kruskal Wallis test; *,P < 0.05, mean ± SEM of n = 5. **(C)** Wildtype or *Bcl6*^*CD4*^ mice were infected i.p. with HKx31 and treated daily with rapamycin or PBS. Mediastinal lymph nodes were removed 15 days after immunization and analyzed for germinal centers. **(D)** The average number of germinal centers per lymph node is depicted. Data are representative of 1 experiment. Kruskal Wallis test; *, P < 0.05, mean ± SEM of n = 3 or 4. **(E)** Sera from anti-CD40L or isotype-treated mice were taken 28 days after immunization and analyzed by ELISA for IgG or **(F)** IgM antibodies specific for HKx31. Freidman test with Dunn’s multiple comparison test; *, P < 0.05, **, P< 0.01, ***, P< 0.001, ****, P< 0.0001, mean ± SEM of n = 7 to 10. Data are representative of two independent experiments. **(G)** Sera from wildtype or *Bcl6*^*CD4*^ mice were taken 28 days after immunization and analyzed by ELISA for IgG or **(H)** IgM antibodies specific for HKx31. Data are of two experiments. Freidman test with Dunn’s multiple comparison test; *, P < 0.05, **, P< 0.01, ****, P ≤ 0.0001, mean ± SEM of n = 11 to 15.

To determine the impact of reducing germinal center formation on the responding antibody response, we analyzed sera for HKx31-specific IgG and IgM, 28 days following infection. Mice given anti-CD40L antibody had reduced HKx31-specific IgG, but HKx31-specific IgM levels were maintained compared to isotype controls (Fig. 1, E and F). Similarly, in *Bcl6*^*CD4*^ mice, HKx31-specific IgG, but not IgM, was reduced compared to wildtype controls (Fig. 1, G and H). These data are consistent with previous findings that long-lived IgM, but not IgG, plasma cells develop in the absence of germinal center formation (24). Interestingly, although germinal centers were reduced in rapamycin-treated mice compared to controls, the amount of HKx31-specific IgG was similar between rapamycin and PBS-treated mice (Fig. 1, E and G). Moreover, as we previously reported, rapamycin-treated mice had more HKx31-specific IgM compared to control mice. However, the increase in HKx31-specific IgM was not observed in *Bcl6*^*CD4*^ mice or mice treated with anti-CD40L compared to controls (Fig. 1, F and H). It is intriguing that germinal centers were reduced comparably in rapamycin-treated, anti-CD40L-treated, and *Bcl6*^*CD4*^ mice, yet only the rapamycin-treated mice displayed increased levels of HKx31-specific IgM, without a reduction in IgG. These data suggest that the increased HKx31-specific IgM in rapamycin-treated mice is not simply due to a reduction in germinal centers and reduced class switching to IgG, implying that rapamycin impacts the generation of influenza-specific IgM by a different mechanism. Although germinal centers were typically regarded as the major site of antibody isotype switching, several groups demonstrated extrafollicular isotype switching (25–28). In fact, Roco et al., recently reported that the majority of isotype switching occurs prior to germinal center formation (29). The fact that IgG is reduced in anti-CD40L-treated and *Bcl6*^*CD4*^ mice, but not in rapamycin-treated mice suggests that anti-CD40L signals by T_FH_ are required for extrafollicular isotype switching, and that rapamycin is modulating a distinct signaling pathway. Together, these results indicate effective methods to limit germinal center formation following influenza infection, and interestingly, these methods have distinct impacts on the antibody response. As germinal center-independent IgM memory cells can be long-lived, these cells may be good candidates to target for vaccine development (24, 30–33).

### Germinal center inhibition decreases high avidity H3-specific IgG antibodies

To determine whether the synthesis of high avidity antibodies specific for HA was impaired in mice with fewer germinal centers, we analyzed sera from mice harvested 28 days after HKx31 infection by an H3 HA ELISA with increasing concentrations of urea. Similar to our observations with whole virus, H3-specific IgG levels were reduced in anti-CD40L-treated and *Bcl6*^*CD4*^ mice compared to controls (Fig. 2, A and B), while H3-specific IgM levels were maintained (Fig. 2, C and D). This suggests that the immunodominance of the HA epitope in shaping the IgM repertoire can be established independent of germinal center competition. Furthermore, in both models, rapamycin-treated mice had more H3-specific IgM relative to PBS-treated control mice (Fig. 2, C and D). As expected, the avidity of H3-specific IgG antibodies produced in the anti-CD40L-treated mice was significantly reduced compared to isotype control-treated mice, with a similar trend observed in *Bcl6*^*CD4*^ mice relative to wildtype controls (Fig. 2, E and F). These findings are consistent with the established theory that germinal centers are the main site of antigen-driven affinity maturation (11). Interestingly, despite a reduction of germinal centers in mice treated with rapamycin, the level of IgG specific for whole virus was not reduced (Fig. 1, E and G); however, there was a trend in lower amounts of high avidity H3-specific IgG antibodies relative to PBS-treated controls (Fig. 2, E and F). These data demonstrate that class switching to IgG occurs outside germinal centers, and further highlight the role of germinal centers in fine-tuning the avidity of IgG to specific epitopes. In contrast to the IgG antibodies, the absence of germinal centers did not reduce the avidity of H3-specific IgM compared to controls, except for rapamycin-treated *Bcl6*^*CD4*^ mice (Fig. 2, G and H). In general, the avidity of the H3-specific IgM antibodies was reduced in comparison to H3-specific IgG, as indicated by the greater impact of urea on IgM binding compared to IgG antibodies (Fig. 2, E and H). Together, these data indicate that inhibiting germinal center formation, via anti-CD40L or in *Bcl6*^*CD4*^ mice, effectively reduced the prevalence of high avidity influenza-specific IgG antibodies.

**Fig. 2.**
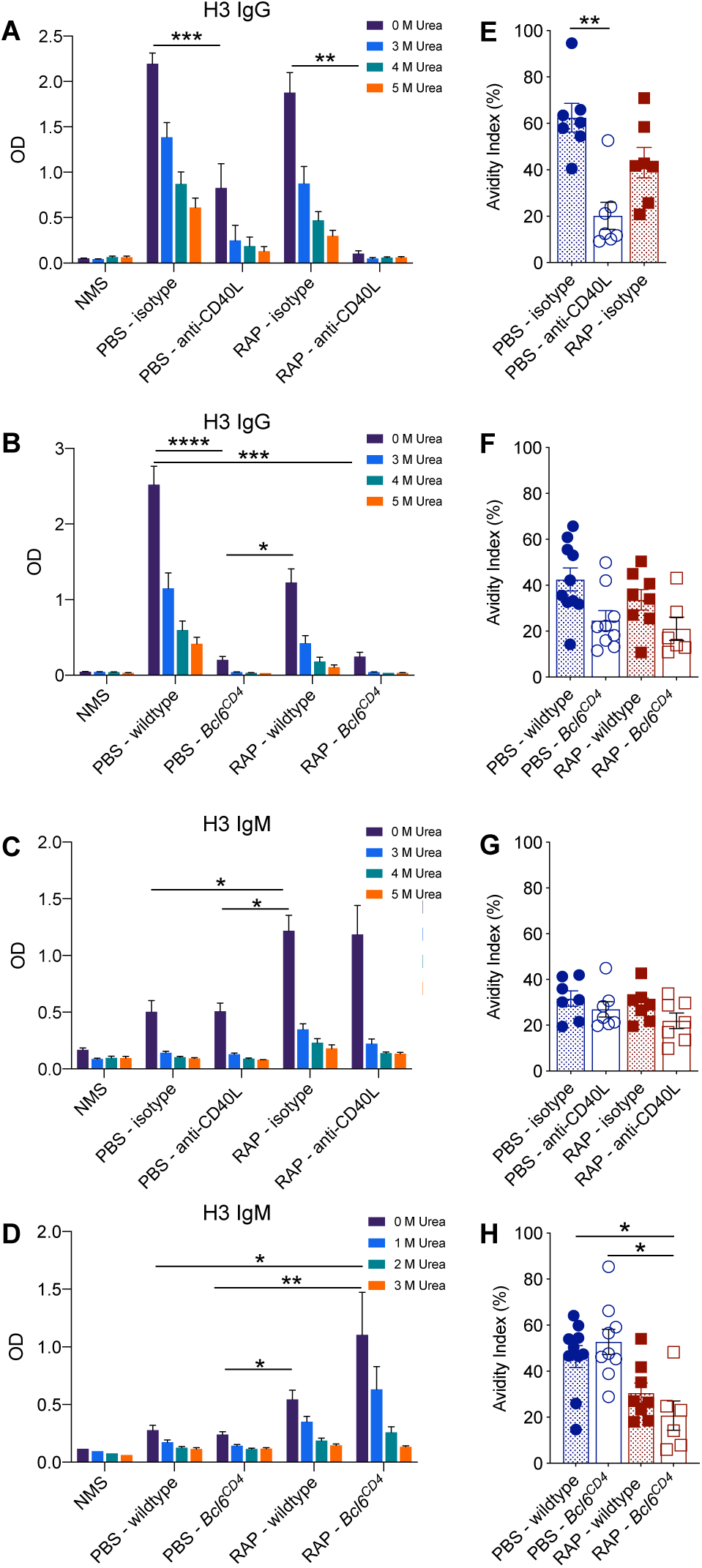
Germinal center inhibition decreases high avidity H3-specific IgG antibodies. Sera from anti-CD40L or isotype-treated mice were taken 28 days after immunization with HKx31 and analyzed by ELISA for **(A)** H3-specific IgG or (**C)** H3-specific IgM. Sera from wildtype or *Bcl6*^*CD4*^ mice were taken 28 days after immunization and analyzed by ELISA for **(B)** H3-specific IgG or (**D)** H3-specific IgM. To determine the avidity of the antibodies, urea was added at decreasing concentrations and the avidity indexes were calculated as the ratio of OD at 3M urea/OD with no urea (Abs 3M urea /AbsDiluent). (**E-F**) Avidity indexes for H3 IgG, and (**G-H**) H3 IgM are depicted. Data are representative of two independent experiments. Kruskal Wallis test; *, P < 0.05, **, P < 0.01, ***, P< 0.001, ****, P ≤ 0.0001, mean ± SEM of n = 5 to 10, NMS and Vn1203 positive control n = 4 to 5.

### Reducing high avidity H3-specific IgG antibodies does not promote broadly reactive influenza antibodies

To determine whether reducing germinal center formation and the frequency of high avidity H3-specific IgG antibodies gives rise to an increase in broadly reactive influenza antibodies, we tested sera from mice infected with HKx31 (H3N2) for antibodies specific for an H5 HA protein from A/Vietnam/1203/04 (Vn1203) or a recombinant H5N1 virus that contains H5 and N1 from Vn1203 in combination with the same internal genes as HKx31 (ΔVn1203). The frequency of broadly reactive, H5-specific IgG antibodies was negligible for all groups of mice (Fig. 3, A and B). IgG antibodies specific for the whole ΔVn1203 virus were detectable in HKx31-infected mice; however, the reduction in high avidity anti-H3 IgG antibodies in mice treated with anti-CD40L did not lead to an increase in broadly reactive IgG antibodies compared to control mice (Fig. 3C). Likewise, *Bcl6*^*CD4*^ mice did not have an increase in broadly reactive IgG compared to wildtype mice (Fig. 3D). Not surprisingly, given the diminished opportunity for germinal center affinity maturation, the avidity of the cross-reactive, ΔVn1203-specific IgG antibodies was reduced in anti-CD40L-treated and *Bcl6*^*CD4*^ mice compared to control mice (Fig. 3, D and F). Although all groups of mice lacked IgG antibodies specific for H5, all HKx31-infected mice had an increase in H5-specific IgM antibodies compared to naïve mice (NMS) (Fig. 3, G and I). Likewise, IgM antibodies specific for the whole ΔVn1203 virus increased in all HKx31-infected mice, relative to uninfected mice (Fig. 3, K and M). There was a trend for increased H5 and ΔVn1203-specific IgM in rapamycin-treated mice compared to PBS controls, but not in mice treated with anti-CD40L (Fig. 3, G and K). However, reducing high avidity antibodies in anti-CD40L-treated or *Bcl6*^*CD4*^ mice did not increase the amount of cross-reactive H5 or ΔVn1203-specific IgM relative to control mice with intact germinal centers (Fig. 3, G-M). Interestingly, rapamycin-treated *Bcl6*^*CD4*^ mice that had a reduction in high avidity H3-specific IgM antibodies (Fig. 2, G and H) also showed an increase in high avidity ΔVn1203-specific IgM antibodies (Fig. 3N). Together, these results indicate that limiting the formation of the high avidity H3-specific IgG antibodies by reducing germinal center formation is not sufficient to increase the prevalence of broadly reactive influenza antibodies. Furthermore, rapamycin alters the antibody repertoire via a mechanism independent of germinal center reduction, and this impact is enhanced in *Bcl6*^*CD4*^ mice that lack T_FH._

**Fig. 3.**
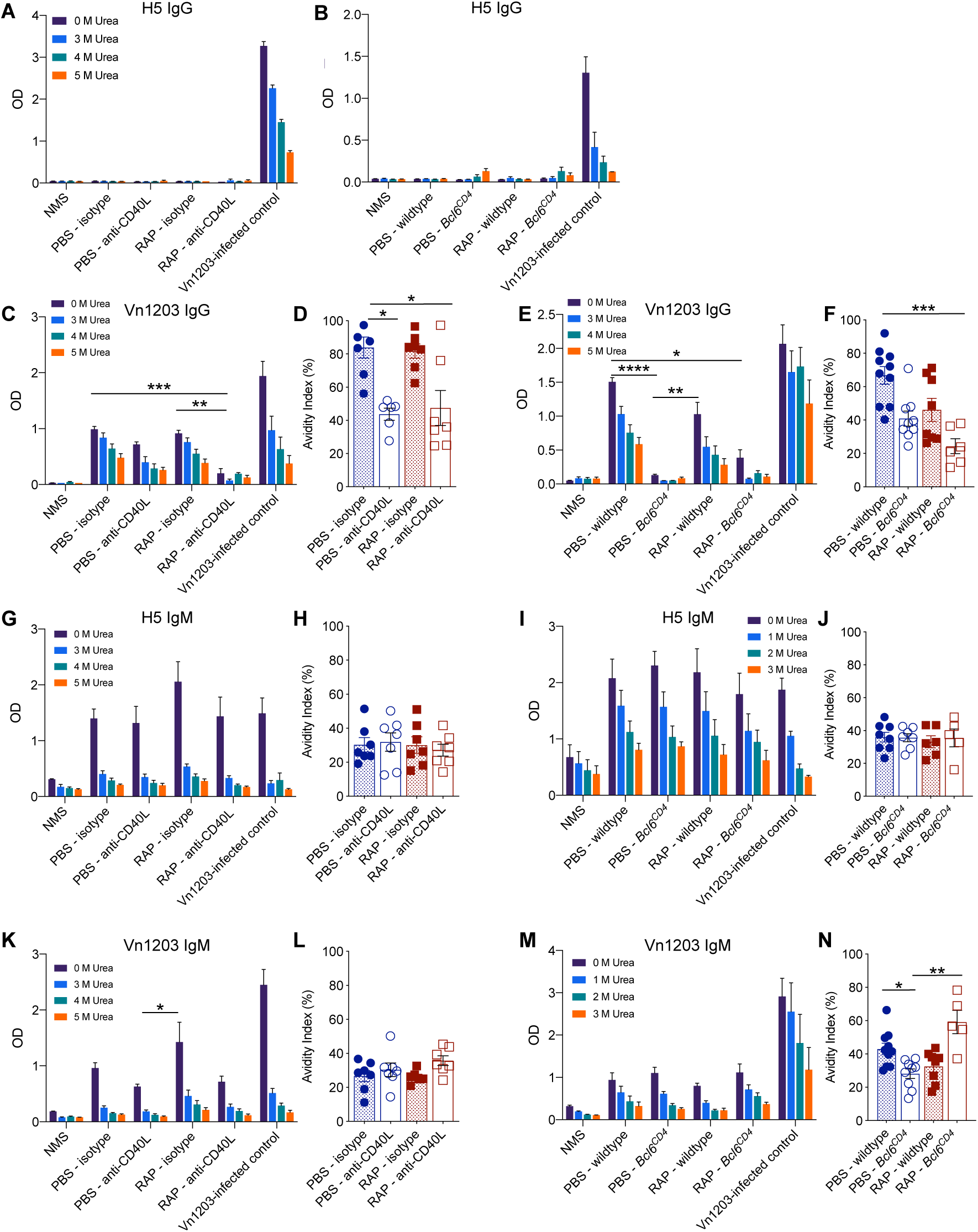
Reducing high avidity H3-specific IgG antibodies does not promote influenza cross-protective antibodies. Sera from (**A**) anti-CD40L or isotype-treated mice, or (**B**) wildtype and *Bcl6*^*CD4*^ mice were taken 28 days after immunization with HKx31 and analyzed by ELISA for (**A-B**) H5-specific IgG (**C-F**) ΔVn1203-specific IgG, (**G-J**) H5-specific IgM, or (**K-N**) ΔVn1203-specific IgM. Avidity indexes were determined with 3M urea **(D**,**F**,**H**,**J**,**L) or 2M urea (N)**. Data are representative of at least two independent experiments. Kruskal Wallis test; *, P < 0.05, **, P < 0.01 ***, P < 0.001, mean ± SEM of n = 5 to 10 for the samples, NMS and Vn1203 positive control n = 3 to 5.

### Limiting high avidity antibodies in intact germinal centers is not sufficient to boost broadly reactive influenza antibodies

Our study highlights that influenza-specific IgM and IgG antibodies can be maintained independent of germinal centers. It is possible that germinal center-independent IgM antibodies are limited by IgG antibodies that are generated outside of the germinal center. Additionally, within germinal centers, broader IgM repertoires may develop in the absence of high avidity antibodies. Therefore, we sought to limit high avidity antibodies while keeping the germinal centers intact. To this end, we infected *Aicda*^-/-^ mice, which cannot undergo somatic hypermutation or isotype class switching, and therefore cannot produce IgG or high avidity antibodies in the germinal centers (34). The *Aicda*^-/-^ and wildtype C57BL/6 mice were infected i.p. with HKx31, and antibodies in the sera were analyzed 28 days later. As expected, *Aicda*^-/-^ mice did not have any HKx31-specific IgG following infection, in both rapamycin and PBS-treated mice (Fig. 4A). In addition, the level of HKx31-specific IgM increased in *Aicda*^-/-^ mice treated with rapamycin or PBS compared to wildtype mice, which likely reflects accumulation of IgM due to a block in class switching to IgG (Fig. 4B). Similar to anti-CD40L-treated or *Bcl6*^*CD4*^ mice lacking germinal centers, *Aicda*^-/-^ mice that lack high avidity IgG did not have an increase in broadly reactive IgM antibodies relative to wildtype mice (Fig. 4C), suggesting that removal of the high avidity IgG antibodies was not sufficient to increase broadly reactive antibodies. Remarkably, *Aicda*^-/-^ mice treated with rapamycin had the highest levels of ΔVn1203-specific IgM antibodies and the highest avidity antibodies compared to all other groups of mice (Fig. 4D), which was similar to rapamycin-treated *Bcl6*^*CD4*^ mice. Together, these data confirm that removing high avidity H3-specific antibodies is not sufficient to promote the generation of broadly reactive influenza antibodies. Additionally, rapamycin enhances the generation of broadly reactive antibodies via a mechanism other than removing competition from high avidity antibodies.

**Fig. 4.**
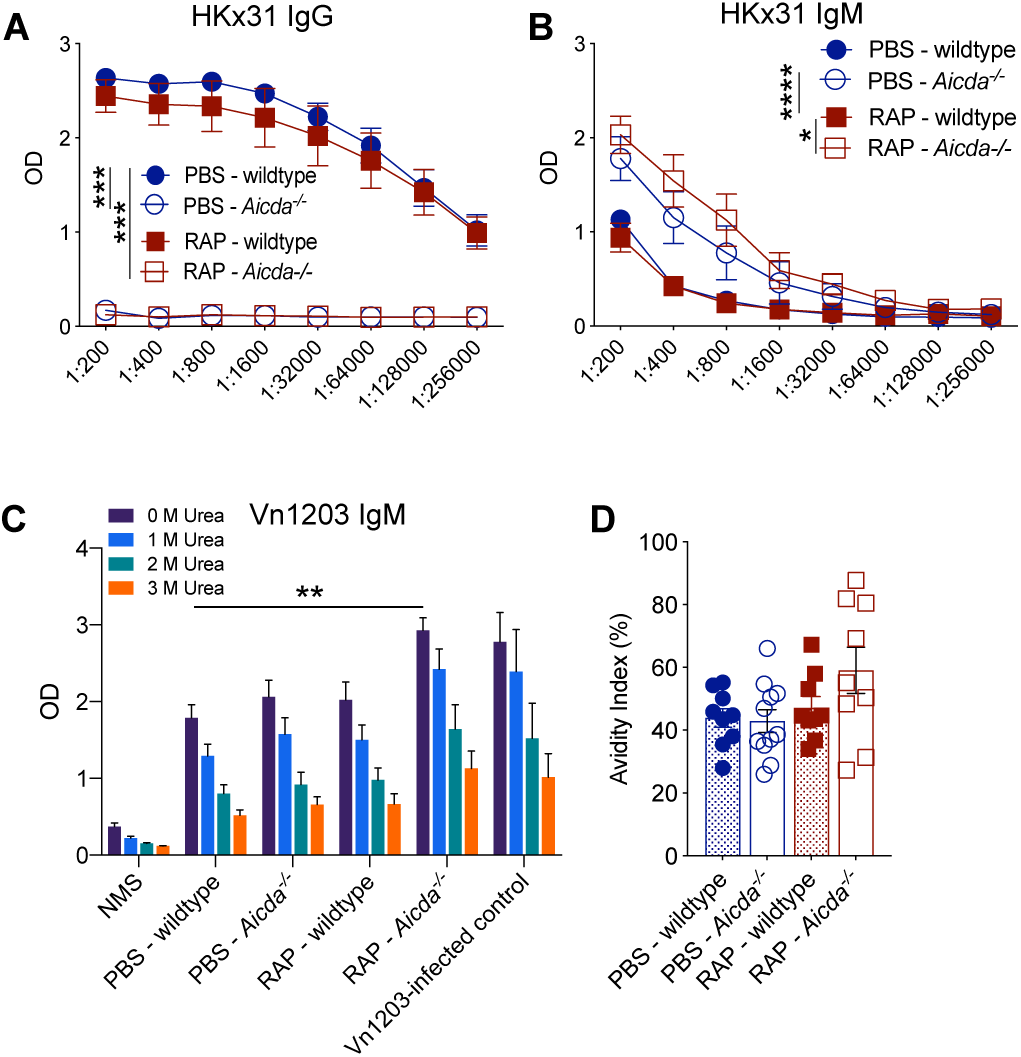
Limiting high avidity antibodies via *Aicda* deletion is not sufficient to boost influenza cross-protective antibodies. Wildtype or *Aicda*^*-/-*^ mice were infected i.p. with HKx31 and treated daily with rapamycin or PBS. Sera were taken 28 days after immunization and analyzed by ELISA for HKx31-specific IgG **(A)** or HKx31-specific IgM (**B**). Freidman test with Dunn’s multiple comparison test; *, P < 0.05, ***, P < 0.001, ****, P< 0.0001, mean ± SEM of n = 6 to 7. Data are representative of one experiment. **(C)** Sera from mice immunized 28 days prior with HKx31 were analyzed for ΔVn1203-specific IgM antibodies with increasing concentrations of urea. Kruskal Wallis test; **, P < 0.01 **(D)** The avidity index was calculated with 3M urea. Data are representative of one experiment. Mean ± SEM of n = 9 to 11. NMS and ΔVn1203-infected control n = 3 to 6.

### Reducing high avidity IgG antibodies is not sufficient to boost immunity to subsequent heterosubtypic influenza infections

Our previous work demonstrated that rapamycin enhanced protection to heterosubtypic influenza infections when given during vaccination with an H3N2 virus (18). This protection was B cell-dependent and transferred to naïve mice via serum, indicating that rapamycin enhanced protection by altering the antibody response. When analyzing the antibody response to the whole virus, there is only a modest increase in the levels of ΔVn1203-specific IgM antibodies in mice treated with rapamycin compared to control mice (Fig. 3). However, we previously demonstrated that rapamycin altered the specificities of both the IgG and IgM antibodies generated during infection, and there were significant differences in antibodies specific for particular epitopes in the rapamycin and PBS-treated mice (18). Thus, although mice that lack high avidity IgG antibodies via germinal center or *Aicda* deletion did not have an increase in the levels of broadly reactive influenza antibodies, altered antibody specificities may enhance heterosubtypic protection. Therefore, we tested whether restricting the production of high avidity IgG antibodies in mice infected with an H3N2 virus impacted survival following a secondary challenge with an H5N1 virus (ΔVn1203). Mice were infected i.p. with HKx31 and treated with either rapamycin or PBS as a control for 28 days. The following day, mice were given an intranasal challenge with ΔVn1203 and monitored for survival and weight loss. As we reported previously, rapamycin treatment significantly increased survival compared to PBS-treated controls (Fig. 5). However, germinal center reduction via anti-CD40L treatment or in *Bcl6*^*CD4*^ mice was not sufficient to enhance protection against a heterosubtypic virus (Fig. 5, A-D). Likewise, deletion of high avidity antibodies in *Aicda*^-/-^ mice did not enhance protection following an H5N1 infection (Fig. 5, E and F). These data indicate that simply reducing germinal center formation or high avidity IgG antibodies is not sufficient to increase broadly reactive influenza immunity. Notably, the mice that received anti-CD40L and *Bcl6*^*CD4*^ mice treated with rapamycin were protected from ΔVn1203 infection at levels similar to rapamycin-treated wildtype mice even though these mice had minimal HKx31-specific or cross-reactive IgG antibodies. This suggests that rapamycin enhances heterosubtypic immunity in a manner independent of germinal center formation and high avidity IgG antibodies. Furthermore, the fact that wildtype, anti-CD40L-treated, and *Bcl6*^*CD4*^ mice given rapamycin had higher levels of HKx31-specific IgM than the PBS-treated cohorts, which was also observed for cross-reactive IgM antibodies, supports the notion that immunity to conserved portions of influenza virus may be enhanced by increasing broadly reactive IgM antibodies. The protective role of IgM antibodies in rapamycin-treated mice is most evident by the survival of *Aicda*^*-/-*^ mice treated with rapamycin, which completely lack IgG antibodies. Further studies to assess the protective capacity of rapamycin through germinal center-independent IgM antibodies may provide valuable insight into enhancing immunity to conserved influenza epitopes. Together, our data show that reducing germinal center formation and limiting the prevalence of high avidity antibodies is not sufficient to enhance heterosubtypic immunity to influenza viruses.

**Figure 5.**
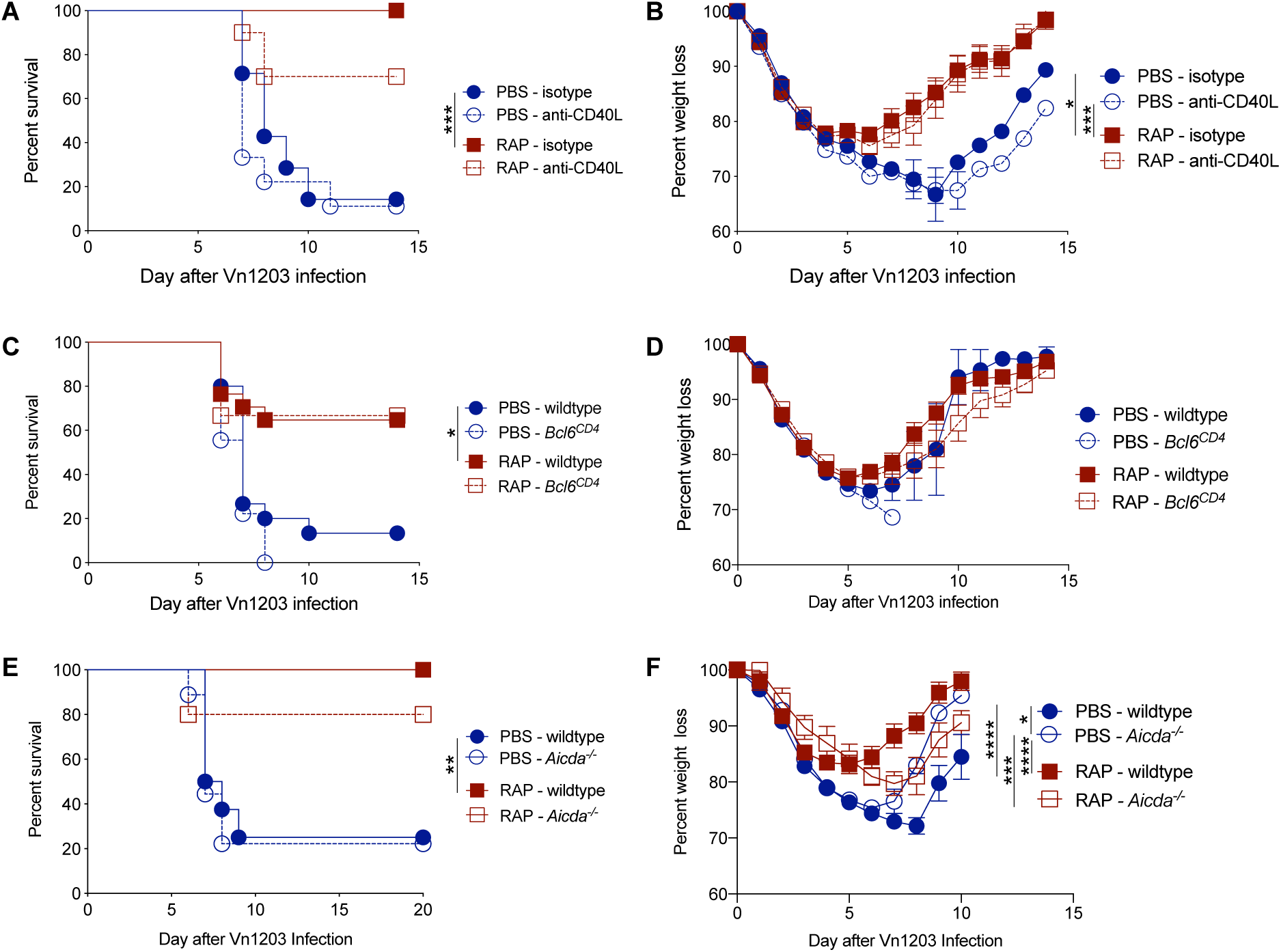
Reducing high avidity IgG antibodies is not sufficient to boost immunity to subsequent heterosubtypic influenza infections. C57BL/6 mice treated with anti-CD40L or isotype control antibody were infected with ΔVn1203 i.n., 30 days after HKx31 immunization and daily injections of PBS or rapamycin. Mice were monitored for **(A)** survival **p<0.01, (Mantel-Cox) and **(B)** weight loss, **p<0.001 (linear mixed-effects model). Data are representative of two independent experiments, (n = 7 to 10). Wildtype and *Bcl6*^*CD4*^ mice were infected with ΔVn1203 i.n., 30 days after HKx31 immunization and daily injections of PBS or rapamycin. Mice (n = 9-17/group) were monitored for **(C)** survival, **p<0.01(Mantel-Cox) and **(D)** weight loss. Data are representative of three independent experiments. C57BL/6 mice and *Aicda*^*-/-*^ mice were infected with ΔVn1203 i.n., 30 days after HKx31 immunization and daily injections of PBS or rapamycin. Mice were monitored for **(E)** survival **p<0.01, (Mantel-Cox) and **(F)** weight loss, *p<0.05 (linear mixed-effects model. Data are representative of two independent experiments, (n = 5 to 9).

In contrast to the general dogma that broadly reactive antibodies are rare due to favored selection of high avidity antibodies in germinal centers, our data demonstrate that restricting development of high avidity IgG antibodies is not sufficient to increase the prevalence of broadly reactive influenza antibodies. The fact that antibodies specific for the conserved regions of influenza did not develop in the absence of high avidity antibodies implies that other factors limit the development of broadly reactive antibodies. The precursor frequency of broadly reactive B cells was recently shown to be similar to B cells specific for the variable region, indicating that precursor frequency does not limit broadly reactive antibodies (17). Moreover, in the absence of the HA variable region, antibodies specific for the conserved stalk are generated, demonstrating the immunogenic potential of these epitopes (1, 17). During the 2009 H1N1 pandemic, antibodies specific for the HA stalk were much more prevalent in the general population relative to prior years, suggesting that steric hindrance of the conserved epitopes on the intact virus does not completely block the generation of broadly reactive antibodies (35). However, infection and vaccination with the 2009 pandemic H1N1 strain was also associated with a higher risk of narcolepsy and Guillain-Barré syndrome (36–42). Further, some antibodies specific for the HA stalk region were shown to be polyreactive (16, 43). Thus, antibodies specific for conserved influenza epitopes may have a higher likelihood of cross-reacting to self-proteins, and therefore their generation may be limited by tolerance mechanisms.

## Materials and Methods

### Mice

Female, 7-8 week-old, C57BL/6J mice and *CD4.Cre* mice were obtained from The Jackson Laboratory. Mice with loxP-flanked *Bcl6* alleles were previously described (21) and were crossed with *CD4.Cre* mice to generate *Bcl6*^*fl/fl*^.*CD4-Cre* (*Bcl6*^*CD4*^) mice that lack *Bcl6* in T cells. As controls, the *Bcl6*^*CD4*^ mice were compared to either *Bcl6*^*fl/fl*^ mice negative for *CD4-Cre*, or *Bcl6*^*+/+*^ mice positive for *CD4-cre. Aicda*^-/-^ mice (34) were generated at St. Jude from frozen sperm obtained from the RIKEN BioResource Center. Female, 7-8 week old, *Aicda*^-/-^ mice were used in experiments with age-matched female C57BL/6J mice. All mice were maintained under specific pathogen free conditions at St. Jude Children’s Research Hospital, and all animal studies were approved by the Institutional Animal Care and Use Committee.

### Virus and Infections

The HKx31 and ΔVn1203 viruses were constructed using the eight-plasmid reverse genetics system (44) and contained the six internal genes of A/Puerto Rico/8/34 (PR8) and genes encoding the HA and NA surface proteins from either A/HKx31 (H3N2) or A/Vietnam/1203/04 (H5N1), respectively. Mice were given 1×10^8^ EID_50_ of HKx31 intraperitoneally (i.p.) diluted in PBS. Beginning one day prior to HKx31 administration, mice received daily i.p. injections of 1.5µg rapamycin (Rapamune; Wyeth) diluted in PBS, or PBS alone as a control. Mice treated with anti-CD40L antibody were given i.p. injections of 200 mg of anti-CD40L, or an isotype control, on day 6 and 8 relative to HKx31 vaccination. For secondary challenge, after four weeks of rapamycin or PBS injections, mice were anesthetized with Avertin (2,2,2-tribromoethanol) and challenged intranasally with 4.5×10^5^ EID_50_ of ΔVn1203. Mice were monitored daily for weight loss and clinical signs of disease. Mice determined to be moribund were euthanized via CO_2_ asphyxiation.

### Germinal Center Analysis

Mice were euthanized on day 15 following HKx31 infection. Mediastinal lymph nodes were removed and fixed in 4% formaldehyde, embedded in paraffin, sectioned, and stained with anti-Bcl6 (sc-858;Santa Cruz) antibodies. Images were acquired with a Nikon TiE microscope equipped with a 10X, 0.3NA objective, motorized stage and DS-Ri2 CMOS camera. Image capture, processing and analysis was performed using NIS Elements software (Nikon Instruments).

### HKx31-Specific ELISA

Microtiter plates (Nunc) were coated with lysed whole HKx31 diluted in PBS at 100ng/well overnight at 4°C. Plates were washed and incubated with serum samples for 2h, then washed and incubated with goat anti-mouse IgG (1030-04;Southern Biotechnology Associates) or goat anti-mouse IgM (1020-04;Southern Biotechnology Associates) for 1hr. The IgG and IgM antibodies were detected using P-nitrophenyl phosphate (Sigma-Aldrich) added for 30 min to 4hrs at 25°C and the OD was measured at 405nm in a microplate reader (Molecular Devices).

### H3-, H5-, and Vn1203-Specific ELISA with Urea

Microtiter plates (Nunc) were coated with H3 or H5 protein at 35ng/well for IgG assays or 70ng/well for IgM assays or lysed whole Vn1203 diluted in PBS at 10ng/well for IgG assays, or 20ng/well for IgM assays, overnight at 4°C. Plates were washed three times with 0.5% Tween in PBS and blocked with 2.5% FBS in PBS for 1h at 25°C on a shaker. Plates were incubated with serum samples for 1h on a shaker and washed three times. Dilutions of urea in PBS (1M-5M) were made and added at 25°C, for 15 minutes, to destabilize antigen-antibody interactions and make a relative comparison of antibody binding avidity. The plates were then washed three times and incubated with either goat anti-mouse IgG Fc-HRP (1033-05;Southern Biotechnology Associates) or goat anti-mouse IgM HRP (1140-05;Southern Biotechnology Associates) for 30 minutes at 25°C on the shaker. Plates were washed three times and TMB (3,3’,5,5’-tetramethylbenzidine) substrate was added in the dark at 25°C for 15 minutes. The reaction was stopped via addition of 1N sulfuric acid. Absorbances were measured at 450 and 550nm in a microplate reader (Molecular Devices). Background signal (550nm) was subtracted from Absorbance (OD450) values to give the relative OD. An avidity index (AI) value was calculated for sera using either 2M or 3M of urea, in accordance with the rate of antigen-antibody destabilization, and compared to no urea, or diluent alone (AbsUREA/AbsDiluent) (45).

### Statistical Analysis

All data were graphed and analyzed using Prism 7.04 software (GraphPad Software), except for analysis of weight-loss as described below. Quantitative differences between more than two groups were compared using the Kruskal-Wallis test followed by a Dunn’s multiple comparison test. ELISA curves with serial dilutions of sera were compared using the Freidman test for repeated measures, followed by a Dunn’s multiple comparison test. Survival experiments were analyzed by the Kaplan-Meier survival probability estimates. Weight-loss comparisons were made using a generalized linear mixed model analysis. Models were fit with the lme4 R package, with individual mice included as a random effect to correct for the non-independence of the data. Residual plots were used to ensure homoscedasticity of the residuals. P values of less than 0.05 were considered significant.

## Acknowledgments

We would like to thank Jeremy C. Crawford for help with statistical analysis, Paul G. Thomas for helpful discussions and reagents, Ashley Castellaw, Carly Lewis, Katherine Anderson, Krishna Patel, and Juan Mejia for technical assistance, the St. Jude Animal Resource Center for their support and excellent animal care, and the St. Jude Cryopreservation Laboratory for rederivation of the *Aicda*^*-/-*^ mice.

## Author contributions

R.K., J.L.J., A.L.D., and M.A.M. designed the experiments. R.K., J.L.J., J.L, D.C.B., and M.A.M. performed the experiments and analyzed data. R.K. and M.A.M. wrote the manuscript. R.K., J.L.J., J.L, D.C.B., A.L.D and M.A.M edited the manuscript. All authors approved the final manuscript.

